# A hypothalamic circuit mechanism underlying the impact of stress on memory and sleep

**DOI:** 10.1101/2024.10.17.618467

**Authors:** Alyssa Wiest, John J. Maurer, Franz Weber, Shinjae Chung

## Abstract

Stress profoundly affects sleep and memory processes. Stress impairs memory consolidation, and similarly, disruptions in sleep compromise memory functions. Yet, the neural circuits underlying stress-induced sleep and memory disturbances are still not fully understood. Here, we show that activation of CRH^PVN^ neurons, similar to acute restraint stress, decreases sleep and impairs memory in a spatial object recognition task. Conversely, inhibiting CRH^PVN^ neurons during stress reverses stress-induced memory deficits while slightly increasing the amount of sleep. We found that both stress and stimulation of CRH^PVN^ neurons activate neurons in the lateral hypothalamus (LH), and that their projections to the LH are critical for mediating stress-induced memory deficits and sleep disruptions. Our results suggest a pivotal role for CRH^PVN^ neuronal pathways in regulating the adverse effects of stress on memory and sleep, an important step towards improving sleep and ameliorating the cognitive deficits that occur in stress-related disorders.

## INTRODUCTION

Stress disrupts memory and sleep (Cheeta et al., 1997; Conrad, 2010; Han et al., 2012; Joëls et al., 2006; Kim and Dimsdale, 2007; Lo Martire et al., 2019; Park et al., 2015; Pawlyk et al., 2008; Roozendaal, 2002; Sandi and Pinelo-Nava, 2007; Schwabe et al., 2022). In particular, spatial memory is susceptible to the effects of stress as well as sleep disturbances (Binder et al., 2012; Graves et al., 2003; Ishikawa et al., 2014; Li et al., 2012; Lopes da Cunha et al., 2019; Palchykova et al., 2006; Prince et al., 2014; Smith and Rose, 1997; Valdivia et al., 2024). Although numerous studies have established links between stress, memory, and sleep, the neural mechanisms underlying the impact of stress on both memory and sleep remain poorly understood.

Corticotropin-releasing hormone (CRH) plays a crucial role in regulating autonomic, endocrine, and behavioral responses to stress (Bale and Vale, 2004; de Kloet et al., 2005; Herman and Tasker, 2016). It has been shown to be involved in regulating spontaneous wakefulness and stress-induced arousal (Chang and Opp, 2004, 2002, 2001, 1999, 1998; Sanford et al., 2015). Furthermore, activation of CRH neurons in the paraventricular nucleus of the hypothalamus (CRH^PVN^) strongly promotes wakefulness and is associated with triggering stress-related behaviors, such as grooming and digging, in the absence of stress (Füzesi et al., 2016; Li et al., 2020; Mitchell et al., 2024; Ono et al., 2020). Optogenetic activation of CRH^PVN^ neurons causes an increase in the release of corticosterone, a primary stress hormone, which further emphasizes the importance of these neurons in regulating physiological responses to stress (Füzesi et al., 2016). The role of CRH^PVN^ neurons in modulating stress responses and sleep-wake patterns highlights their potential impact on cognitive functions. However, the specific neural pathways involved and whether targeted interventions in these circuits can mitigate stress-induced disruptions in memory and sleep are still unknown.

CRH^PVN^ neurons densely project to the lateral hypothalamus (CRH^PVN→LH^), a brain region implicated in the regulation of sleep-wake states and memory (Bonnavion et al., 2016; Füzesi et al., 2016; Graebner et al., 2015; Li et al., 2020; Mitchell et al., 2024; Ono et al., 2020; Rho and Swanson, 1987). Optogenetic activation of CRH^PVN^ neurons has been shown to increase c-Fos labeling in the LH, confirming the functional connection between these two brain regions (Füzesi et al., 2016; Mitchell et al., 2024). This pathway has been studied for its role in modulating wakefulness, stress-induced behavioral changes, and alterations in motivational drive (Füzesi et al., 2016; Li et al., 2020; Mitchell et al., 2024; Ono et al., 2020). Different populations of neurons within the lateral hypothalamus (LH) have been linked to the regulation of memory processes. Optogenetic activation of hypocretin/orexin neurons in the LH after the learning phase of the novel object recognition (NOR) task causes sleep fragmentation and impairs memory (Rolls et al., 2011). Similarly, activating melanin-concentrating hormone (MCH) neurons in the LH significantly impairs memory in the NOR task, underscoring the critical role of these neurons in memory functions (Izawa et al., 2019). Although the CRH^PVN→LH^ pathway has been studied for its role in regulating sleep-wake cycles and stress responses, how this projection influences both sleep and memory following stress is largely unclear, constituting a major gap in our understanding of the neural circuitry involved in the interplay between stress, sleep, and cognitive functions.

By employing acute restraint stress coupled with optogenetic manipulations as well as a spatial object recognition (SOR) task, we investigated the role of CRH^PVN^ neurons and their projections to the LH in regulating memory performance and sleep-wake states following stress.

## RESULTS

### Activation of CRH^PVN^ neurons impairs spatial object recognition memory and decreases sleep

By performing an SOR task combined with electroencephalogram (EEG) and electromyography (EMG) recordings, we tested whether activation of CRH^PVN^ neurons following training is sufficient to impair spatial memory performance and reduce sleep (**Fig. 1**). To investigate this, we optogenetically activated CRH^PVN^ neurons using the stabilized step-function opsin (SSFO), a double mutant excitatory channelrhodopsin. CRH-Cre mice were injected with an AAV encoding Cre-inducible SSFO (AAV2-Ef1ɑ-DIO-hChR2-eYFP) or eYFP (AAV2-Ef1ɑ-DIO-eYFP) in the PVN, followed by implantation of an optic fiber above the injection site and electrodes for EEG/EMG recordings (**Fig. 1A**). Two weeks later, we performed the SOR task (**Fig. 1B**). During habituation (ZT1-1.5, days 1 and 2), mice were familiarized with the open field where the training and testing would occur for two 5-minute sessions per day. A visual cue was attached to one wall to help orient the mice within the field. On the training day (ZT1-1.5, day 3), mice were placed in the open field with two identical objects (small glass bottles) for three 5-minute training sessions. Immediately following training, optogenetic stimulation was performed for 1 hour (1 second step pulses at 120-240 second intervals, 4 mW, ZT1.5-2.5) followed by EEG and EMG recordings (ZT2.5-7.5). Twenty-four hours later, during the test session (ZT1-1.5, day 4), mice were placed in the open field with the two familiar objects, one of which was moved to a new location, for one 5-minute session. We found that eYFP control mice showed a significant increase in preference for the moved object and a positive discrimination ratio during the test session (**Figs. 1C-E**; paired t-test, test vs. training, p = 0.016). Conversely, SSFO mice exhibited no discernible preference between the moved and familiar objects, indicating impaired performance in this memory task (**Fig. 1D**; paired t-test, test vs. training, p = 0.895). Accordingly, the discrimination ratio of the SSFO mice was significantly lower than that of the eYFP control mice (**Fig. 1E**; unpaired t-test, p = 0.007), suggesting that the activation of CRH^PVN^ neurons after training impairs memory performance.

**FIGURE 1.**
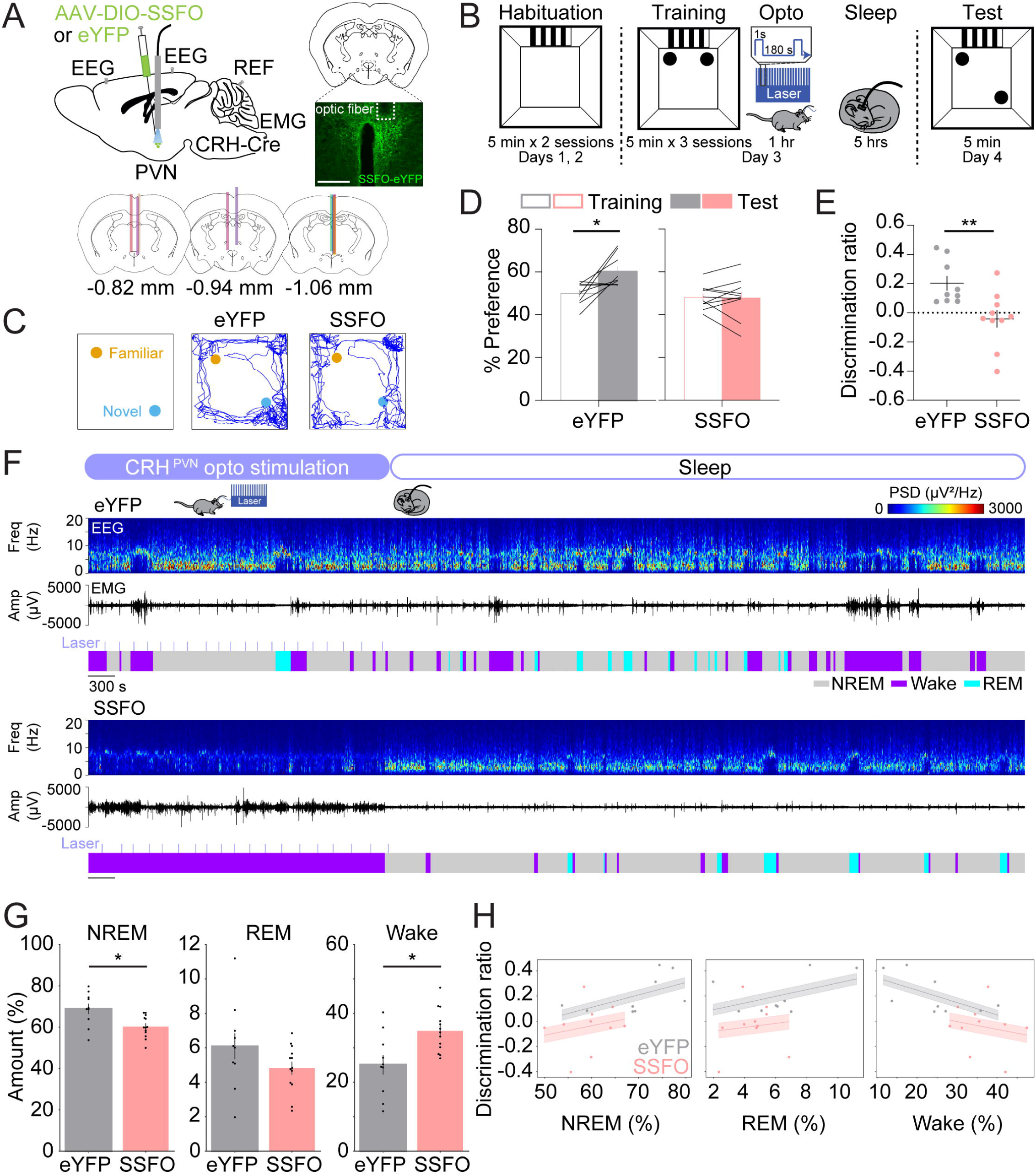
Activation of CRH^PVN^ neurons impairs spatial object recognition memory and decreases sleep. (**A**) Top left, schematic of optogenetic activation experiments with simultaneous EEG and EMG recordings. Mouse brain figure adapted from the Allen Reference Atlas – Mouse Brain (https://atlas.brain-map.org). Top right, fluorescence image of the PVN in a CRH-Cre mouse injected with AAV2-Ef1α-DIO-hChR2-eYFP into the PVN. Scale bar, 300 µm. Bottom, location of optic fiber tracts. **(B)** Schematic indicating the timing of the spatial object recognition task in eYFP and SSFO mice. **(C)** Representative locomotor trajectory line graphs, based on nose x– and y-coordinates, of eYFP and SSFO mice during the test session. **(D)** Preference (%) for the moved object during training and testing sessions in eYFP (gray, left) and SSFO (pink, right) mice. Outline-only bars indicate the training day, while filled bars represent the test day. n = 9 eYFP mice and 10 SSFO mice. **(E)** Discrimination ratio during the testing session for eYFP (gray, left) and SSFO (pink, right) mice. n = 9 eYFP mice and 10 SSFO mice. **(F)** Example recording of an eYFP (top) and SSFO mouse (bottom) during 1 hour laser stimulation, laser indicated by the light blue lines, and sleep for 2 hours following laser stimulation. Shown are EEG power spectra, EMG amplitude, and color-coded brain states. **(G)** Percentage of time spent in NREM sleep, REM sleep, and wakefulness for the combined 3-hour period immediately following training on the memory task for eYFP and SSFO mice. n = 10 eYFP mice and 13 SSFO mice. **(H)** Linear mixed model analysis illustrating the relationship between discrimination ratio (y-axis) and percentages of NREM, REM, and wake states (x-axis) in eYFP and SSFO mice. Each panel represents a different sleep state. Grey dots indicate individual data points for eYFP mice (n = 9), and pink dots for SSFO mice (n = 10). Lines represent fitted values from the linear mixed effects model. Error clouds represent the standard error of the mean from residuals. Bars, averages across mice; dots and lines, individual mice; error bars, s.e.m. Unpaired and paired t-tests, **P < 0.01; *P < 0.05.

In addition, we examined how the activation of CRH^PVN^ neurons after training in the SOR task impacts sleep. Several studies have highlighted critical windows for memory consolidation, suggesting that sleep disturbances or the manipulation of neural activity within the first 4-6 hours after the final training session most severely impair memory consolidation (Bayer and Bertoglio, 2020; Graves et al., 2003; Hong et al., 2024; Palchykova et al., 2006; Prince et al., 2014; Smith and Rose, 1996; Smith et al., 1998). We found that the amount of wakefulness was significantly increased and NREM sleep was reduced during the first 3 hours of the sleep recording following training (including the 1-hour optogenetic stimulation interval and the following 2-hour sleep sessions; **Figs. 1F-G, S1A-B**; unpaired t-tests, NREM: p = 0.011, REM: p = 0.172, wake: p = 0.011), but not during the last 3 hours (**Fig. S1C**). In the first hour, optogenetic activation of CRH^PVN^ neurons significantly increased wakefulness while decreasing NREM and REM sleep (**Fig. S1A**, unpaired t-test, NREM: p = 2.700e-5, REM: p = 0.017, Wake: p = 1.900e-5). During the following 2-hour sleep session (after laser stimulation ended), NREM sleep was increased (**Fig. S1B**, unpaired t-test, NREM: p = 0.019). Next, we investigated whether changes in sleep-wake states in the first 3 hours were correlated with memory performance. The correlation between the percentage of wakefulness and the discrimination ratio was marginally significant, suggesting that the overall increase in wakefulness and decrease in sleep may contribute to the impaired performance in the SOR task (**Fig. 1H**; Linear Mixed Model, wake percentage: z = –1.716, p = 0.086, group comparison: z = –0.572, p = 0.567).

### Inhibition of CRH^PVN^ neurons during stress improves spatial object recognition memory and sleep

Next, we investigated whether the activity of CRH^PVN^ neurons contributes to the memory impairment and disturbed sleep observed after stress. To first assess the impact of stress on the consolidation of spatial memory, we subjected mice to 1 hour of acute restraint stress immediately after the final training session in the SOR task (**Fig. 2A**). Control mice, which received no manipulations, exhibited a significant increase in their preference for the moved object and a positive discrimination ratio (**Figs. 2B-D**; paired t-test, test vs. training, p = 0.030). In contrast, mice subjected to acute restraint stress showed no preference for the moved object and a discrimination ratio close to zero (**Figs. 2B-D**; paired t-test, test vs. training, p = 0.568). The discrimination ratio was significantly lower in restraint stress mice compared with control mice (**Fig. 2D**; unpaired t-test, p = 0.046).

**FIGURE 2.**
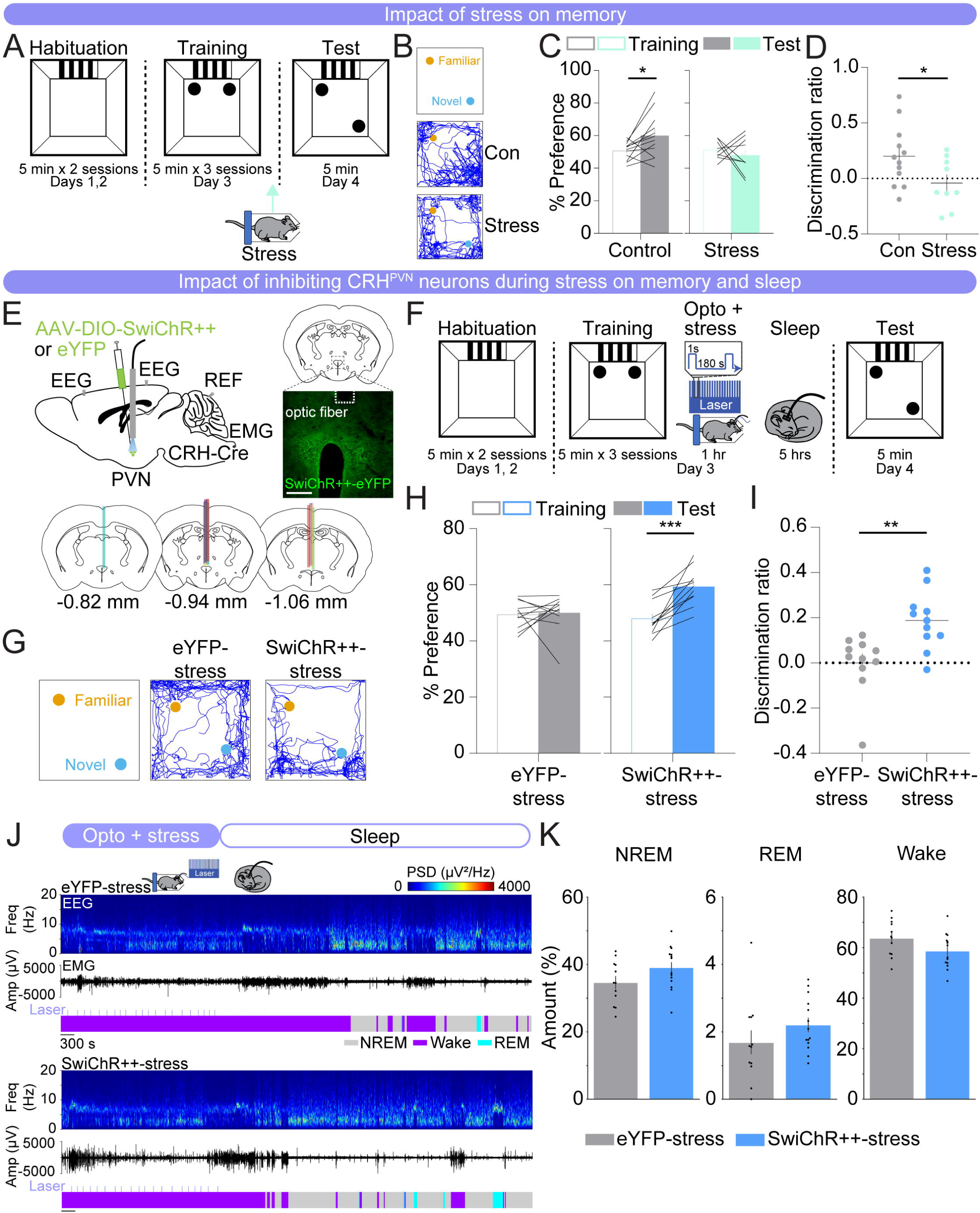
Inhibition of CRH^PVN^ neurons during stress improves spatial object recognition memory and sleep. (**A**) Schematic indicating the timing of the spatial object recognition task in no restraint stress (control) and restraint stress (stress) mice. Arrow indicates the timing of restraint stress. **(B)** Representative locomotor trajectory line graphs, based on nose x– and y-coordinates, of control and stress mice during the test session. **(C)** Preference (%) for the moved object during training and testing sessions in control (gray, left) and restraint stress (mint, right) mice. Outline-only bars indicate the training day, while filled bars represent the test day. n = 12 control mice and 9 restraint stress mice. **(D)** Discrimination ratio during the testing session for control (gray, left) and restraint stress (mint, right) mice. n = 12 control mice and 9 restraint stress mice. **(E)** Top left, schematic of optogenetic inhibition experiments with simultaneous EEG and EMG recordings. Top right, fluorescence image of the PVN in a CRH-Cre mouse injected with AAV2-Ef1α-DIO-SwiChR++-eYFP into the PVN. Scale bar, 300 µm. Bottom, location of optic fiber tracts. **(F)** Schematic indicating the timing of the SOR task in eYFP-stress and SwiChR++-stress mice. **(G)** Representative locomotor trajectory line graphs, based on nose x– and y-coordinates, of eYFP-stress and SwiChR++-stress mice during the test session. **(H)** Preference (%) for the moved object during training and testing sessions in eYFP-stress (gray, left) and SwiChR++-stress (blue, right) mice. Outline-only bars indicate the training day, while filled bars represent the test day. n = 11 eYFP-stress mice and 11 SwiChR++-stress mice. **(I)** Discrimination ratio during the testing session for eYFP-stress (gray, left) and SwiChR++-stress (blue, right) mice. n = 11 eYFP-stress mice and 11 SwiChR++-stress mice. **(J)** Example recording of an eYFP-stress (top) and SwiChR++-stress mouse (bottom) during 1 hour laser stimulation, laser indicated by the light blue lines, during restraint stress and sleep for 2 hours following laser stimulation. Shown are EEG power spectra, EMG amplitude, and color-coded brain states. **(K)** Percentage of time spent in NREM sleep, REM sleep, and wakefulness for the combined 3-hour period immediately following training on the SOR task for eYFP-stress and SwiChR++-stress mice. n = 12 eYFP-stress mice and 14 SwiChR++-stress mice. Bars, averages across mice; dots and lines, individual mice; error bars, s.e.m. Unpaired and paired t-tests, ***P < 0.001; **P < 0.01; *P < 0.05.

Subsequently, we tested whether optogenetic inhibition of CRH^PVN^ neurons during restraint stress is sufficient to reverse the memory performance deficits and sleep disturbances caused by acute restraint stress. CRH-Cre mice were bilaterally injected with an AAV encoding the Cre-inducible bistable chloride channel, SwiChR++ (AAV2-EF1ɑ-DIO-SwiChR++-eYFP) or eYFP (AAV2-EF1ɑ-DIO-eYFP) in the PVN, followed by optic fiber implantation above the injection sites (**Fig. 2E**) (Berndt et al., 2016). Immediately following training, CRH^PVN^ neurons in these mice were optogenetically inhibited during the 1 hour of restraint stress (1 second step pulses at 120-240 second intervals, 4 mW, ZT1.5-2.5) and their sleep-wake states were recorded afterwards (ZT2.5-7.5) (**Fig. 2F**). We found that eYFP-stress mice exhibited no discernible preference between the moved and familiar objects on the test day, indicating impaired performance in the memory task (**Figs. 2G-I**; paired t-test, test vs. training, p = 0.831). In contrast, SwiChR++-mediated inhibition of CRH^PVN^ neurons during stress resulted in a significant preference for the moved object (**Figs. 2G-I**; paired t-test, test vs. training, p = 3.000e-4). The discrimination ratio of the SwiChR++-stress mice was significantly higher than that of the eYFP-stress mice, indicating improved memory task performance (**Fig. 2I**; unpaired t-test, p = 0.003). This suggests that the activity of CRH^PVN^ neurons during restraint stress following training contributes to the observed memory impairment.

We then examined whether inhibiting CRH^PVN^ neurons during restraint stress impacts sleep-wake states (**Figs. 2J, K**). During restraint stress, both eYFP– and SwiChR++-stress mice were awake for the entire 1-hour restraint stress period (**Fig. S2B**). When averaging the entire 3-hour period following optogenetic inhibition during restraint stress, SwiChR++-stress mice had overall slightly more NREM and REM sleep and less wakefulness compared with eYFP-stress mice (**Fig. 2K**; unpaired t-tests, p = 0.089, 0.222, 0.080 for NREM, REM, and wake, respectively). The percentage of each brain state and the discrimination ratio were not significantly correlated (**Fig. S2A**). These results indicate that although inhibiting CRH^PVN^ neurons during stress improves memory and sleep, the increase in sleep may not directly contribute to the memory improvement. This suggests that CRH^PVN^ neuron activity during stress may influence sleep and memory through independent processes.

### Inhibition of CRH^PVN^ neurons does not affect spatial object recognition memory and sleep

Next, we tested whether inhibiting CRH^PVN^ neurons alone affects memory and sleep. CRH-Cre mice were bilaterally injected with AAV2-EF1ɑ-DIO-SwiChR++-eYFP or AAV2-EF1ɑ-DIO-eYFP in the PVN, followed by optic fiber implantation (**Fig. 3A**). Immediately following training, both eYFP and SwiChR++ mice received laser stimulation (1 second step pulses at 120-240 second intervals, 4 mW, ZT1.5-2.5). We found that both groups exhibited an increased preference for the moved object on the test day (**Figs. 3C-D**; paired t-tests, test vs. training, eYFP: p = 0.011, SwiChR++: p = 0.048). The discrimination ratio of the SwiChR++ mice did not significantly differ from that of the eYFP mice (**Fig. 3E**; unpaired t-test, p = 0.373). Both groups of mice spent similar amounts of time in NREM sleep, REM sleep, and wakefulness (**Figs. 3F-G**; unpaired t-tests, NREM: p = 0.536, REM: p = 0.579, wake: p = 0.490). This indicates that the optogenetic inhibition of CRH^PVN^ neurons alone does not significantly alter spatial memory task performance or sleep-wake states (**Figs. 3C-G**).

**FIGURE 3.**
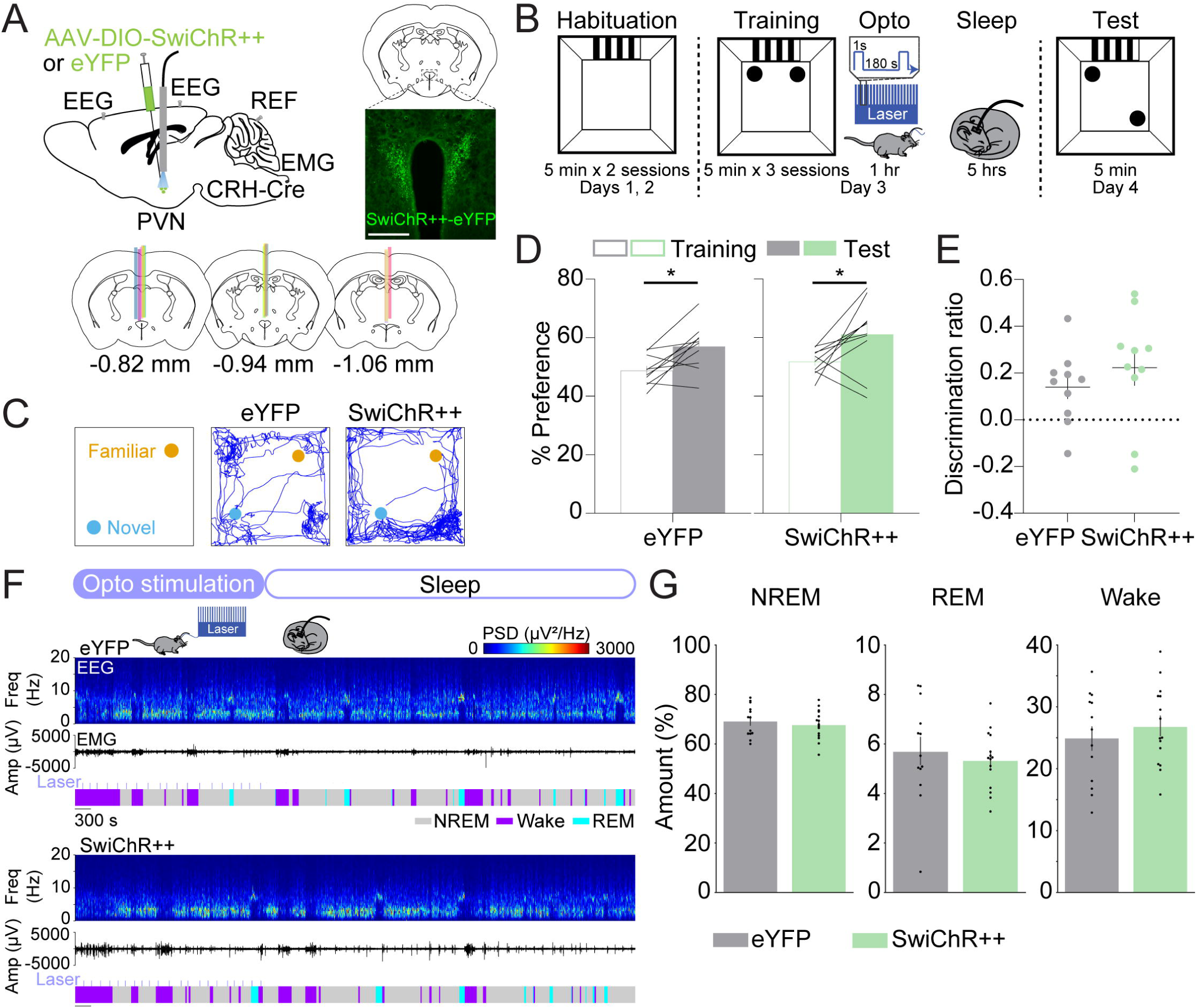
Inhibition of CRH^PVN^ neurons does not affect spatial object recognition memory and sleep. (**A**) Top left, schematic of optogenetic inhibition experiments with simultaneous EEG and EMG recordings. Top right, fluorescence image of the PVN in a CRH-Cre mouse injected with AAV2-Ef1α– DIO-SwiChR++-eYFP into the PVN. Scale bar, 400 µm. Bottom, location of optic fiber tracts. **(B)** Schematic indicating the timing of the SOR task in eYFP and SwiChR++ mice. **(C)** Representative locomotor trajectory line graphs, based on nose x– and y-coordinates, of eYFP and SwiChR++ mice during the test session. **(D)** Preference (%) for the moved object during training and testing sessions in eYFP (gray, left) and SwiChR++ (green, right) mice. Outline-only bars indicate the training day, while filled bars represent the test day. n = 10 eYFP mice and 10 SwiChR++ mice. **(E)** Discrimination ratio during the testing session for eYFP (gray, left) and SwiChR++ (green, right) mice. n = 10 eYFP mice and 10 SwiChR++ mice. **(F)** Example recording of an eYFP (top) and SwiChR++ mouse (bottom) during 1 hour laser stimulation, laser indicated by the light blue lines, and sleep for 2 hours following laser stimulation. Shown are EEG power spectra, EMG amplitude, and color-coded brain states. **(G)** Percentage of time spent in NREM sleep, REM sleep, and wakefulness for the combined 3-hour period immediately following training on the SOR task for eYFP and SwiChR++ mice. n = 13 eYFP mice and 15 SwiChR++ mice. Bars, averages across mice; dots and lines, individual mice; error bars, s.e.m. Unpaired and paired t-tests, ***P < 0.001; **P < 0.01; *P < 0.05.

### Inhibition of CRH^PVN→LH^ projections during restraint stress improves spatial object recognition memory and sleep

CRH^PVN^ neurons densely project to the LH, a brain region implicated in the regulation of stress, sleep-wake states, and memory (Bonnavion et al., 2016; Füzesi et al., 2016; Graebner et al., 2015; Li et al., 2020; Mitchell et al., 2024; Ono et al., 2020; Rho and Swanson, 1987). We found that both CRH^PVN^ activation and restraint stress increased the number of c-Fos+ cells in the LH, suggesting that the LH is a downstream target that could be involved in mediating the effects of CRH^PVN^ neuron activation and stress on memory and sleep (**Figs. 4A-F**).

**FIGURE 4.**
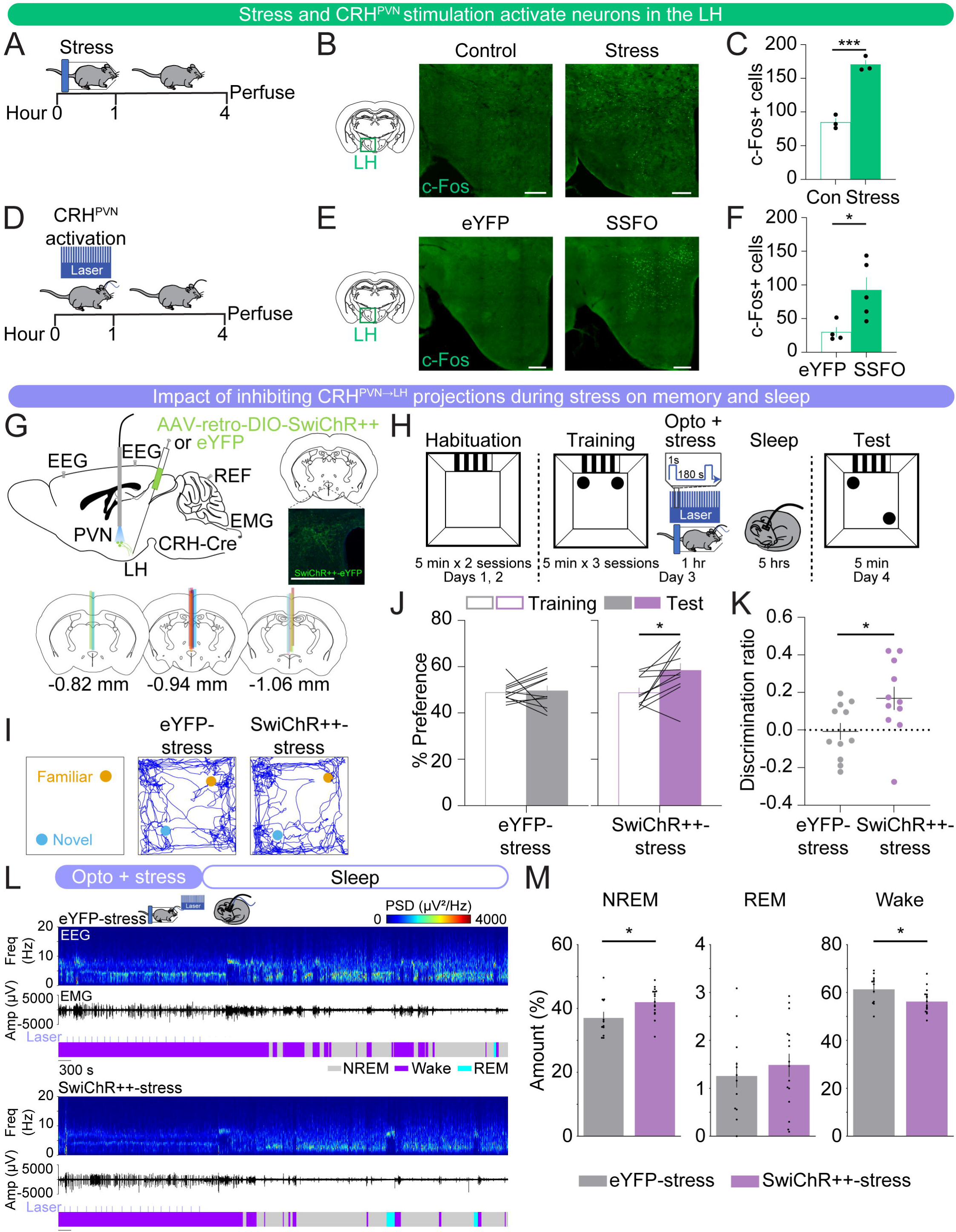
Inhibition of CRH^PVN→LH^ projections during restraint stress improves spatial object recognition memory and sleep. (**A**) Schematic indicating the timing of restraint stress and perfusion for c-Fos experiments. **(B)** Representative images of c-Fos expression (green) in the lateral hypothalamus of no restraint stress (control) and restraint stress (stress) mice. Scale bar, 300 µm. **(C)** Bar graphs indicating the number of c-Fos positive cells, bilateral, counted in the LH averaged from three brain sections/mouse for control (green outline-only bars, left) and restraint stress (green filled bars, right) mice. n = 3 control mice and 3 restraint stress mice. **(D)** Schematic indicating the timing of CRH^PVN^ neuron activation (SSFO) and perfusion for c-Fos experiments. **(E)** Representative images of c-Fos expression (green) in the LH of eYFP control and CRH^PVN^ activation (SSFO) mice. Scale bar, 300 µm. **(F)** Bar graphs indicating the number of c-Fos positive cells, unilateral (due to unilateral virus expression), counted in the LH averaged from two brain sections/mouse for eYFP control (green outline-only bars, left) and SSFO (green filled bars, right) mice. n = 4 eYFP mice and 5 SSFO mice. **(G)** Top left, schematic of optogenetic inhibition experiments with simultaneous EEG and EMG recordings. Top right, fluorescence image of the PVN in a CRH-Cre mouse injected with rAAV2-retro-Ef1α-DIO-SwiChR++-eYFP into the LH. Scale bar, 300 µm. Bottom, location of optic fiber tracts. **(H)** Schematic indicating the timing of the SOR task in retro-eYFP-stress and retro-SwiChR++-stress mice. **(I)** Representative locomotor trajectory line graphs, based on nose x– and y-coordinates, of retro-eYFP-stress and retro-SwiChR++-stress mice during the test session. **(J)** Preference (%) for the moved object during training and testing sessions in retro-eYFP-stress (gray, left) and retro-SwiChR++-stress (purple, right) mice. Outline-only bars indicate the training day, while filled bars represent the test day. n = 11 retro-eYFP-stress mice and 11 retro-SwiChR++-stress mice. **(K)** Discrimination ratio during the testing session for retro-eYFP-stress (gray, left) and retro-SwiChR++-stress (purple, right) mice. n = 11 retro-eYFP-stress mice and 11 retro-SwiChR++-stress mice. **(L)** Example recording of a retro-eYFP-stress (top) and retro-SwiChR++-stress mouse (bottom) during 1 hour laser stimulation, laser indicated by the light blue lines, during restraint stress and sleep for 2 hours following laser stimulation. Shown are EEG power spectra, EMG amplitude, and color-coded brain states. **(M)** Percentage of time spent in NREM sleep, REM sleep, and wakefulness for the combined 3-hour period immediately following training on the SOR task for retro-eYFP-stress and retro-SwiChR++-stress mice. n = 12 retro-eYFP mice-stress and 16 retro-SwiChR++-stress mice. Bars, averages across mice; dots and lines, individual mice; error bars, s.e.m. Unpaired and paired t-tests, ***P < 0.001; **P < 0.01; *P < 0.05.

We tested whether inhibiting CRH^PVN→LH^ projections during restraint stress reversed the memory deficits and sleep disturbances observed after stress. CRH-Cre mice were bilaterally injected with retrograde AAVs encoding Cre-inducible SwiChR++ (rAAV2-retro-Ef1α-DIO-SwiChR++-eYFP) or eYFP (rAAV2-retro-Ef1α-DIO-eYFP) in the LH, followed by optic fiber implantation above the PVN (**Fig. 4G**). Immediately following training, we optogenetically inhibited CRH^PVN→LH^ projections during 1 hour of restraint stress (1 second step pulses at 120-240 second intervals, 4 mW, ZT1.5-2.5), and recorded sleep-wake states afterwards (ZT2.5-7.5) (**Fig. 4H**). Twenty-four hours later, we found that retro-eYFP-stress mice exhibited no discernible preference between the moved and familiar objects on the test day, indicating impaired memory performance after stress (**Figs. 4I-K**; paired t-tests, p = 0.788). In contrast, inhibition of CRH^PVN→LH^ projections during stress immediately following training resulted in a significant increase in the preference for the moved object (**Figs. 4I-K**; paired t-tests, p = 0.014). The discrimination ratio of the retro-SwiChR++-stress mice was significantly higher than that of the retro-eYFP-stress mice, indicating an improved memory task performance (**Fig. 4K**; unpaired t-test, p = 0.031). This suggests that the activity of CRH^PVN→LH^ projections during restraint stress contributes to the memory impairments observed after stress.

Next, we investigated whether inhibiting CRH^PVN→LH^ projections during stress impacts sleep-wake states (**Figs. 4L-M**). During the restraint stress combined with optogenetic manipulation, both retro-eYFP-stress and retro-SwiChR++-stress mice were awake for the entire 1-hour period (**Fig. S3B**). When averaging the entire 3-hour period following optogenetic inhibition during restraint stress, retro-SwiChR++-stress mice had significantly more NREM sleep and spent less time awake compared with retro-eYFP-stress mice (**Fig. 4M**; unpaired t-tests, NREM: p = 0.027, REM: p = 0.513, wake: p = 0.025). The percentage of each brain state and the discrimination ratio were not significantly correlated (**Fig. S3A**), suggesting that the increased sleep following inhibition of CRH^PVN→LH^ projections during stress may not contribute to improved memory. Thus, similar to the effects observed with CRH^PVN^ cell body inhibition during stress, their projections to the LH may regulate sleep and memory through independent mechanisms.

Taken together, these results suggest that the effects of CRH^PVN^ neuronal stimulation on impairments in spatial memory and sleep are, at least in part, mediated by their projections to the LH.

## DISCUSSION

Our study provides crucial insights into how CRH^PVN^ neurons influence spatial memory and sleep-wake patterns in mice after stress. Using optogenetic manipulations, we demonstrated that activating CRH^PVN^ neurons impaired memory in the spatial object recognition task, mirroring the effects of acute restraint stress (**Fig. 1**, **Figs. 2A-D**). In contrast, optogenetic inhibition of CRH^PVN^ neurons during restraint stress improved memory and slightly increased NREM and REM sleep compared with eYFP-stress mice after stress (**Fig. 2**). Inhibition of CRH^PVN^ neurons alone did not have a significant effect on SOR task performance or sleep-wake states (**Fig. 3**). Using c-Fos staining, we found that acute restraint stress and the stimulation of CRH^PVN^ neurons activated neurons in the LH, suggesting that this region is a downstream target of CRH^PVN^ neurons (**Fig. 4**). Optogenetic inhibition of CRH^PVN→LH^ projections during restraint stress improved spatial memory task performance and significantly increased NREM sleep and reduced wakefulness compared with retro-eYFP-stress mice following stress (**Fig. 4**). Our results indicate an important role for CRH^PVN^ neurons in regulating stress-induced cognitive deficits and sleep disturbances, partially through their projections to the LH.

The activation of CRH^PVN^ neurons is a key component of the body’s response to stress. Our study reveals that the activation of CRH^PVN^ neurons significantly increases wakefulness, consistent with prior studies demonstrating that optogenetic activation of CRH^PVN^ neurons promotes wakefulness and triggers stress-associated behaviors, like grooming and digging, in the absence of stress (Füzesi et al., 2016; Li et al., 2020; Mitchell et al., 2024; Ono et al., 2020). Furthermore, our results indicate that the activation of CRH^PVN^ neurons immediately following training negatively impacts performance on a spatial object memory task, and this effect may be partially mediated by their projections to the LH. CRH^PVN^ neurons express mRNA for vesicular glutamate transporter 2 (VGluT2) as a substrate for fast synaptic transmission (Hrabovszky et al., 2005). Therefore, the memory and sleep effects caused by manipulating CRH^PVN→LH^ projections may rely on excitatory glutamatergic projections to neurons in the LH in addition to the release of the CRH peptide. A subset of CRH^PVN^ neurons send collaterals to the median eminence and to the LH, suggesting that the release of corticosterone could partially contribute to sleep and cognitive processes after activation of CRH^PVN^ neurons (Füzesi et al., 2016). Moreover, genetic ablation of LH hypocretin/orexin neurons or *crh* gene knockdown significantly shortens the latency to NREM sleep onset following stress, suggesting a critical role for CRH and LH hypocretin/orexin neurons in mediating not only stress-induced sleep disturbances but also memory deficits (Li et al., 2020).

The LH contains a variety of neurons expressing distinct molecular markers (Mickelsen et al., 2019). GABAergic neurons (Herrera et al., 2016; Venner et al., 2016), glutamatergic neurons (Smith et al., 2024; Wang et al., 2021), neurotensin (Naganuma et al., 2019), and CamKIIɑ-expressing neurons (Heiss et al., 2024) promote wakefulness. In the LH, the activation of MCH neurons after the training period significantly impairs NOR memory (Izawa et al., 2019). Activation of hypocretin/orexin neurons in the LH promotes the release of plasma corticosterone (Bonnavion et al., 2015) and results in sleep fragmentation and impaired memory in the NOR task (Rolls et al., 2011). Given that hypocretin/orexin labels only a subset of LH neurons that become activated after CRH^PVN^ stimulation, it remains to be studied which types of neurons in the LH are innervated by CRH^PVN^ projections and activated by stress in order to mediate the stress-induced impairments in memory and sleep (Mitchell et al., 2024).

Stress impairs hippocampal functions and circuits (Kim et al., 2012, 2007; Tomar et al., 2021; Tomar and McHugh, 2022). In particular, stress decreases the stability of firing rates of place cells in the hippocampus, accompanied by impairments of spatial memory consolidation (Kim et al., 2007). Similarly, amygdala stimulation in rats alters the spatial correlation of place maps and increases variability in the firing rate, whereas corticosterone injection does not (Kim et al., 2012). This suggests that the enhanced amygdalar activity, but not the elevated level of corticosterone, leads to a destabilization of spatial representations within the hippocampus by modifying the firing rates of place cells, mirroring the effects typically seen after behavioral stress (Kim et al., 2012). Hippocampal cells exhibit highly coordinated activity during sleep and resting states. In particular, sharp-wave ripples play a key role in memory consolidation (Buzsáki, 1989; Fernández-Ruiz et al., 2019; Girardeau et al., 2009; Girardeau and Zugaro, 2011; Ramadan et al., 2009). In addition, neuromodulators such as acetylcholine and oxytocin are released in the hippocampus in a brain state-dependent manner, suggesting their vital role in regulating brain state dependent memory functions (Zhang et al., 2024). We found that activation of CRH^PVN^ neurons increased wakefulness, and this increase was correlated with impaired memory. However, this relationship weakened in the context of restraint stress. During stress, inhibition of CRH^PVN^ neurons improves both memory and sleep, while the extent of memory performance was not significantly correlated with the amount of sleep, suggesting that the effects of CRH^PVN^ neurons and their projections to the LH in regulating memory and sleep after stress are mediated by separate pathways. Given that hippocampal pathways are crucial for regulating spatial memory, improved memory may be mediated by postsynaptic LH cells projecting to the hippocampal circuits whereas sleep regulation might rely on sleep-regulatory neurons in the hypothalamus and brainstem, which are not directly connected to the hippocampal circuits (Bittencourt, 2011; Bittencourt et al., 1992; Izawa et al., 2019; Marcus et al., 2001; Peyron et al., 1998). Future studies could explore other factors disrupted by stress, such as sleep-related electrophysiological features and neurotransmitter dynamics, to better understand their contributions to sleep and memory impairments. This approach may offer deeper insights into the complex interplay between stress, memory, and sleep.

While we observed that 1 hour of CRH^PVN^ inhibition during the light phase in the SOR task did not significantly alter sleep-wake states, another study found that chemogenetic inhibition of these neurons during the dark phase led to a reduction in spontaneous wakefulness (Ono et al., 2020). Given that the activity of CRH^PVN^ neurons is under circadian influence, it remains to be investigated whether their activity differentially modulates sleep and memory processes depending on the circadian cycle, especially after stress.

Together, our findings elucidate the crucial role of CRH^PVN^ neurons and their projections to the LH in modulating the impact of stress on memory and sleep-wake states. Cognitive impairments and sleep disturbances are hallmark features in psychiatric disorders such as post-traumatic stress disorder and major depressive disorder, often manifesting before clinical diagnosis (Chang et al., 1997; Ford and Kamerow, 1989; Koren et al., 2002; Meerlo et al., 2015; Neckelmann et al., 2007; Pearson et al., 2023; Spoormaker and Montgomery, 2008). By identifying the specific contributions of CRH^PVN^ neurons to these dysfunctions, our study provides a foundation for targeted interventions that could mitigate such impairments. Therapeutic strategies that specifically modulate this pathway could not only improve cognitive and sleep outcomes but could also potentially delay the progression of related psychiatric conditions, offering advancements in treating stress-related disorders.

## MATERIALS AND METHODS

### Mice

All experimental procedures were approved by the Institutional Animal Care and Use Committee (IACUC reference # 806197) at the University of Pennsylvania and conducted in compliance with the National Institutes of Health Office of Laboratory Animal Welfare Policy. Experiments were performed in male CRH-IRES-Cre mice (#012704, Jackson Laboratory) or C57BL/6J mice (#000664, Jackson Laboratory) aged 6–18 weeks, weighing 18–25 g at the time of surgery. Mice were randomly assigned to experimental and control groups. Animals were group-housed with littermates on a 12-hour light/12-hour dark cycle (lights on 7 am and off 7 pm) with ad libitum access to food and water.

### Surgical procedures

Mice were anesthetized with isoflurane (1–4%) during the surgery, and placed on a stereotaxic frame (Kopf) while being on a heating pad to maintain body temperature. The skin was incised and small holes were drilled for virus injections and implantations of optic fibers and EEG/EMG electrodes. Two stainless steel screws were inserted into the skull 1.5 mm from midline and 1.5 mm anterior to the bregma, and 2.5 mm from midline and 2.5 mm posterior to the bregma. The reference screw was inserted on top of the cerebellum. Two EMG electrodes were inserted into the neck musculature. Insulated leads from the EEG and EMG electrodes were soldered to a 2 × 3 pin header, which was secured to the skull using dental cement. After surgery, mice were monitored for any signs of pain or distress until fully recovered from anesthesia and ambulatory.

For optogenetic activation experiments (**Fig. 1**), AAV2-Ef1α-DIO-hChR2-eYFP (for activation group) or AAV2-Ef1α-DIO-eYFP (for control group) was unilaterally injected into the PVN (300 nl, AP –0.7 mm; ML ±0.4 mm; DV –4.6 to 4.7 mm from the cortical surface) and an optic fiber (200 μm diameter) was implanted above the injection site in the PVN (AP –0.7 mm; ML ±0.3; DV –4.5 mm).

For optogenetic inhibition experiments (**Figs. 2-3**), AAV2-Ef1α-DIO-SwiChR++-eYFP (for the inhibition group) or AAV2-Ef1α-DIO-eYFP (for the control group) was bilaterally injected into the PVN (200 nl) and an optic fiber was implanted above the middle of the PVN (AP –0.7 mm; ML ±0.00 mm; DV –4 mm).

For retrograde optogenetic inhibition experiments (**Fig. 4**), rAAV2-retro-Ef1α-DIO-SwiChR++-eYFP (for inhibition group) or rAAV2-retro-Ef1α-DIO-eYFP (for control group) was bilaterally injected into the LH (300 nl, AP –1.2 mm; ML ±1 mm; DV –5 to –5.2 mm from the cortical surface) and an optic fiber was implanted above the middle of the PVN (AP –0.7 mm; ML ±0.00 mm; DV –4 mm).

After surgery, mice were allowed to recover for at least 2-3 weeks before beginning experiments.

### Histology

Mice were deeply anesthetized and transcardially perfused with phosphate-buffered saline (PBS) followed by 4% paraformaldehyde (PFA) in PBS. Brains were fixed overnight in 4% PFA and then transferred to a 30% sucrose in PBS solution for at least one night. Brains were embedded and mounted with Tissue-Tek OCT compound (Tissue-Tek, Sakura Finetek) and frozen. 40-60 μm sections were cut using a cryostat (Thermo Scientific HM525 NX) and mounted onto glass slides. Brain sections were washed with PBS followed by counterstaining with Hoechst solution (#33342, Thermo Scientific). Slides were cover-slipped with Fluoromount-G (Southern Biotechnic) and imaged using a fluorescence microscope (Microscope, Leica DM6B; Camera, Leica DFC7000GT; LED, Leica CTR6 LED).

### Immunohistochemistry

For GFP staining to visualize virus expression (**Figs. 1, 4**), brain sections were washed in PBS for 5 minutes, permeabilized using PBST (0.3% Triton X-100 in PBS) for 30 minutes, and incubated in blocking solution (5% normal donkey serum in 0.3% PBST; 017-000-001, Jackson ImmunoResearch Laboratories) for 1 hour. Brain sections were incubated with chicken anti-GFP antibody (1:1000; GFP8794984, Aves Lab) in the blocking solution overnight at 4°C. The following morning, sections were washed in PBS and incubated at room temperature for 2 hours with the donkey anti-chicken secondary antibody conjugated to A594 (1:500; 703-585-155, Jackson ImmunoResearch Laboratories). Afterward, sections were washed with PBS followed by counterstaining with Hoechst solution (#33342, Thermo Scientific). Slides were coverslipped with mounting medium (Fluoromount-G, Southern Biotechnic) and imaged using a fluorescence microscope to verify virus expression and optic fiber placement.

Animals were excluded if no virus expression was detected, or the virus expression/optic fiber tips were not properly localized to the targeted area.

For c-Fos staining (**Fig. 4**), brain sections were washed in PBS for 5 minutes, permeabilized using PBST (0.3% Triton X-100 in PBS) for 30 minutes, and incubated in blocking solution (5% normal donkey serum in 0.3% PBST; 017-000-001, Jackson ImmunoResearch Laboratories) for 1 hour. Brain sections were incubated with rabbit anti-c-Fos antibody (1:1000; 2250S, Cell Signaling Technology) in the blocking solution for 2 days at 4°C. Following primary antibody incubation, sections were washed in PBS and incubated at room temperature for 2 hours with the donkey anti-rabbit secondary antibody conjugated to A594 (1:500; A21207, Thermo Scientific). Afterward, sections were washed with PBS followed by counterstaining with Hoechst solution (#33342, Thermo Scientific). Slides were coverslipped with mounting medium (Fluoromount-G, Southern Biotechnic) and imaged using a fluorescence microscope. The c-Fos positive cells were counted from sections containing the LH.

### Sleep recordings

Sleep recordings were carried out in the animal’s home cage or in a recording cage to which the mouse had been habituated. EEG and EMG electrodes were connected to flexible recording cables via a mini-connector. EEG and EMG signals were recorded using an RHD2132 amplifier (Intan Technologies, sampling rate 1 kHz) connected to the RHD USB Interface Board (Intan Technologies). EEG and EMG signals were referenced to a ground screw placed on top of the cerebellum. To determine the sleep-wake state of the animal, we first computed the EEG and EMG spectrogram for sliding, half-overlapping 5 second windows, resulting in 2.5 second time resolution. To estimate within each 5 second window the power spectral density (PSD), we performed Welch’s method with Hanning window using sliding, half-overlapping 2 second intervals. Next, we computed the time-dependent δ (0.5 to 4 Hz), θ (5 to 12 Hz), σ (12 to 20 Hz) and high γ (100 to 150 Hz) power by integrating the EEG power in the corresponding ranges within the EEG spectrogram. We also calculated the ratio of the θ and δ power (θ/δ) and the EMG power in the range 50 to 500 Hz. For each power band, we used its temporal mean to separate it into a low and high part (except for the EMG and θ/δ ratio, where we used the mean plus one standard deviation as threshold). REMs was defined by a high θ/δ ratio, low EMG, and low δ power. A state was set as NREMs if the δ power was high, the θ/δ ratio was low, and EMG power was low. In addition, states with low EMG power, low δ, but high σ power were scored as NREMs. Wake encompassed states with low δ power and high EMG power and each state with high γ power (if not otherwise classified as REMs). Our algorithm has been published and has a 90.256% accuracy compared with manual scoring by expert annotators (Antila et al., 2022; Chung et al., 2017; Maurer et al., 2024; Schott et al., 2023; Smith et al., 2024; Stucynski et al., 2022; Weber et al., 2018, 2015). We manually verified the automatic classification using a graphical user interface visualizing the raw EEG and EMG signals, EEG spectrograms, EMG amplitudes, and the hypnogram to correct for errors, by visiting each single 2.5 second epoch in the hypnograms. The software for automatic sleep-wake state classification and manual scoring was programmed in Python (available at https://github.com/tortugar/Lab/tree/master/PySleep).

Mice exhibiting two faulty channels of the same type (EEG or EMG) were excluded from sleep analysis due to the challenges in accurately determining sleep states.

### Optogenetic manipulation

Light pulses (1 second step pulses, 4 mW) for SSFO and SwiChR++ experiments were generated by a blue laser (473 nm, Laserglow) and sent through the optic fiber (200 μm diameter, ThorLabs) that connects to the ferrule on the mouse’s head. The timing of laser stimulation for optogenetic activation was randomized within a range of intervals (120-240 second intervals). This laser stimulation protocol was rationally designed based on previous studies (Berndt et al., 2016; Iyer et al., 2016; Selimbeyoglu et al., 2017; Stucynski et al., 2022; Wiegert et al., 2017). TTL pulses to trigger the laser were controlled using a raspberry pi, which was controlled by a custom user interface programmed in Python. Optogenetic manipulations were conducted during the light period for 1 hour. Sleep recordings with stress were performed once to ensure that experimental mice were exposed to acute stress one time.

### Spatial object recognition task

Mice were habituated to being handled twice daily for ∼1-2 minutes per handling for five days prior to beginning the memory task. Mice were also habituated to their recording cages during this time. During the open field habituation session (ZT1-1.5, days 1 and 2), mice were habituated to the training context (13”x13” open field arena) for two 5-minute sessions. A visual cue (a rectangle containing alternating black and white stripes) was attached to one wall of the open field to help the mice orient themselves within the open field. During the training session (ZT1-1.5, day 3), mice were placed in the arena with two identical objects (two small glass bottles) for three 5-minute training sessions. Immediately following the training session, mice were returned to their recording cages. In **Figs. 1-4**, optogenetic manipulation was performed for 1 hour in their recording cage with (**Figs. 2, 4**) or without (**Fig. 1, 3**) restraint stress (ZT1.5-2.5). During the test session (ZT1-1.5, day 4), mice were placed in the arena with the two familiar objects, one displaced to a new location, for one 5-minute test session. Exploration of the objects was defined as the amount of time the mouse had its nose oriented toward the object, within ∼2 cm of the object. Grooming near the object was not counted as exploration (Leger et al., 2013).

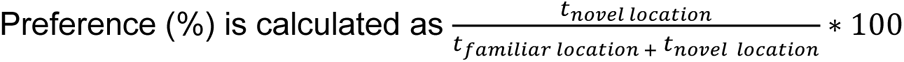

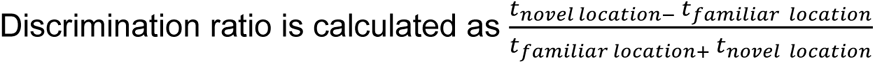

Mice that explored for less than two seconds for two or more training trials or showed a strong preference during the training trials (<40% or >60% preference) were excluded from spatial object recognition analysis. These mice were included in sleep recording analysis. Representative raw traces of locomotion trajectories were plotted using DeepLabCut.

### Locomotion Trajectory

For body part tracking, we used DeepLabCut (version 2.3.9) (Mathis et al., 2018; Nath et al., 2019). Specifically, we labeled 20 frames/video taken from 37 SOR training session videos, then 95% was used for training the model. We used a ResNet-50-based neural network with default parameters for 100,000 training iterations (He et al., 2016; Insafutdinov et al., 2016). This network was then used to analyze videos from similar experimental settings during the SOR test session. We generated representative trajectory line traces based on the nose x– and y-coordinates generated by DeepLabCut. Coordinates which were outside of the arena (i.e. the nose was obscured and couldn’t be tracked) were excluded from the final traces.

### Statistical tests

Statistical analyses were performed using the python modules (scipy.stats, https://scipy.org; pingouin, https://pingouin-stats.org; statsmodels, https://www.statsmodels.org/) and Prism v10.3.1 (GraphPad Software Inc). We did not predetermine sample sizes, but cohorts were similarly sized as in other relevant sleep and memory studies (Izawa et al., 2019; Rolls et al., 2011). All data collection was randomized and counterbalanced. All data are reported as mean + s.e.m. A (corrected) p-value < 0.05 was considered statistically significant for all comparisons. Data were compared using unpaired t-tests, paired t-tests, and a linear mixed model. Statistical results and parameters (exact value of n and what n represents) are presented in the Table S1, figure legends, and results.

## DATA SHARING PLANS

The code used for data analysis is publicly available under: https://github.com/tortugar/Lab. All the data are available from the corresponding author upon reasonable request.

## ACKNOWLEDGMENTS

We thank the members of the Chung and Weber labs for their helpful discussion. This work was funded by the National Institute of Mental Health (R01-MH-136491), the Whitehall Foundation, the Alfred P. Sloan Foundation, and the NIH individual F31 fellowship from the National Heart, Lung, and Blood Institute (F31HL160451, to A.W.).

## AUTHOR CONTRIBUTIONS

Conceptualization: A.W., and S.C.; Methodology: A.W., J.J.M., and F.W.; Software: A.W. and F.W.; Validation: A.W.; Formal Analysis: A.W.; Investigation: A.W.; Data Curation: A.W.; Writing-Original Draft: A.W.; Writing-Review and Editing: A.W., F.W., and S.C.; Visualization: A.W.; Supervision: S.C.; Funding Acquisition: A.W. and S.C.

## FIGURE LEGENDS

**Figure S1.** Effect of CRH^PVN^ activation on sleep-wake states, related to Figure 1. (**A**) Percentage of time spent in NREM sleep, REM sleep, and wakefulness for the 1-hour period of CRH^PVN^ stimulation after training on the SOR memory task for eYFP (gray, left) and SSFO (pink, right) mice. n = 10 eYFP mice and 13 SSFO mice. **(B)** Percentage of time spent in NREM sleep, REM sleep, and wakefulness for the 2-hour period immediately following CRH^PVN^ stimulation for eYFP (gray, left) and SSFO (pink, right) mice. n = 10 eYFP mice and 13 SSFO mice. **(C)** Percentage of time spent in NREM sleep, REM sleep, and wakefulness during the 3-to-6-hour period following CRH^PVN^ stimulation for eYFP (gray, left) and SSFO (pink, right) mice. n = 9 eYFP mice and 9 SSFO mice. One eYFP mouse and four SSFO mice from (A) and (B) were excluded due to having shorter (5 hour long) sleep recordings. Bars, averages across mice; dots and lines, individual mice; error bars, s.e.m. Unpaired and paired t-tests, ***P < 0.001; **P < 0.01; *P < 0.05.

**Figure S2.** Effect of CRH^PVN^ inhibition during restraint stress on sleep-wake states, related to Figure 2. (**A**) Linear mixed model analysis illustrating the relationship between percentages of NREM, REM, and wake states (x-axis) and discrimination ratio (y-axis) in eYFP-stress and SwiChR++-stress mice. Each panel represents a different sleep state. Grey dots indicate individual data points for eYFP-stress mice (n = 10), and blue dots for SwiChR++-stress mice (n = 11). Lines represent fitted values from the linear mixed effects model. Error clouds represent the standard error of the mean from residuals. **(B)** Percentage of time spent in wakefulness for the 1-hour period of CRH^PVN^ inhibition during stress after training on the SOR memory task for eYFP-stress (gray, left) and SwiChR++-stress (blue, right) mice. n = 12 eYFP-stress mice and 14 SwiChR++-stress mice. All mice in both groups spent the entire 1-hour stress period awake. Bars, averages across mice; dots and lines, individual mice; error bars, s.e.m. Unpaired and paired t-tests, ***P < 0.001; **P < 0.01; *P < 0.05.

**Figure S3.** Effect of CRH^PVN→LH^ inhibition during restraint stress on sleep-wake states, related to Figure 4. (**A**) Linear mixed model analysis illustrating the relationship between percentages of NREM, REM, and wake states (x-axis) and discrimination ratio (y-axis) in retro-eYFP-stress and retro-SwiChR++-stress mice. Each panel represents a different sleep state. Grey dots indicate individual data points for retro-eYFP-stress mice (n = 11), and purple dots for retro-SwiChR++-stress mice (n = 11). Lines represent fitted values from the linear mixed effects model. Error clouds represent the standard error of the mean from residuals. **(B)** Percentage of time spent in wakefulness for the 1-hour period of CRH^PVN→LH^ inhibition during stress after training on the SOR memory task for retro-eYFP-stress (gray, left) and retro-SwiChR++-stress (purple, right) mice. n = 12 retro-eYFP-stress mice and 16 retro-SwiChR++-stress mice. All mice in both groups spent the entire 1-hour stress period awake. Bars, averages across mice; dots and lines, individual mice; error bars, s.e.m. Unpaired and paired t-tests, ***P < 0.001; **P < 0.01; *P < 0.05.

**Table S1. Detailed statistical analysis of all datasets, related to Figures 1-4 and S1-3**.

## Notes

### Competing Interest Statement

The authors have declared no competing interest.

