## Supplemental Figures 1-3 and Table S1 for "A hypothalamic circuit mechanism underlying the impact of stress on memory and sleep"

Figure S1

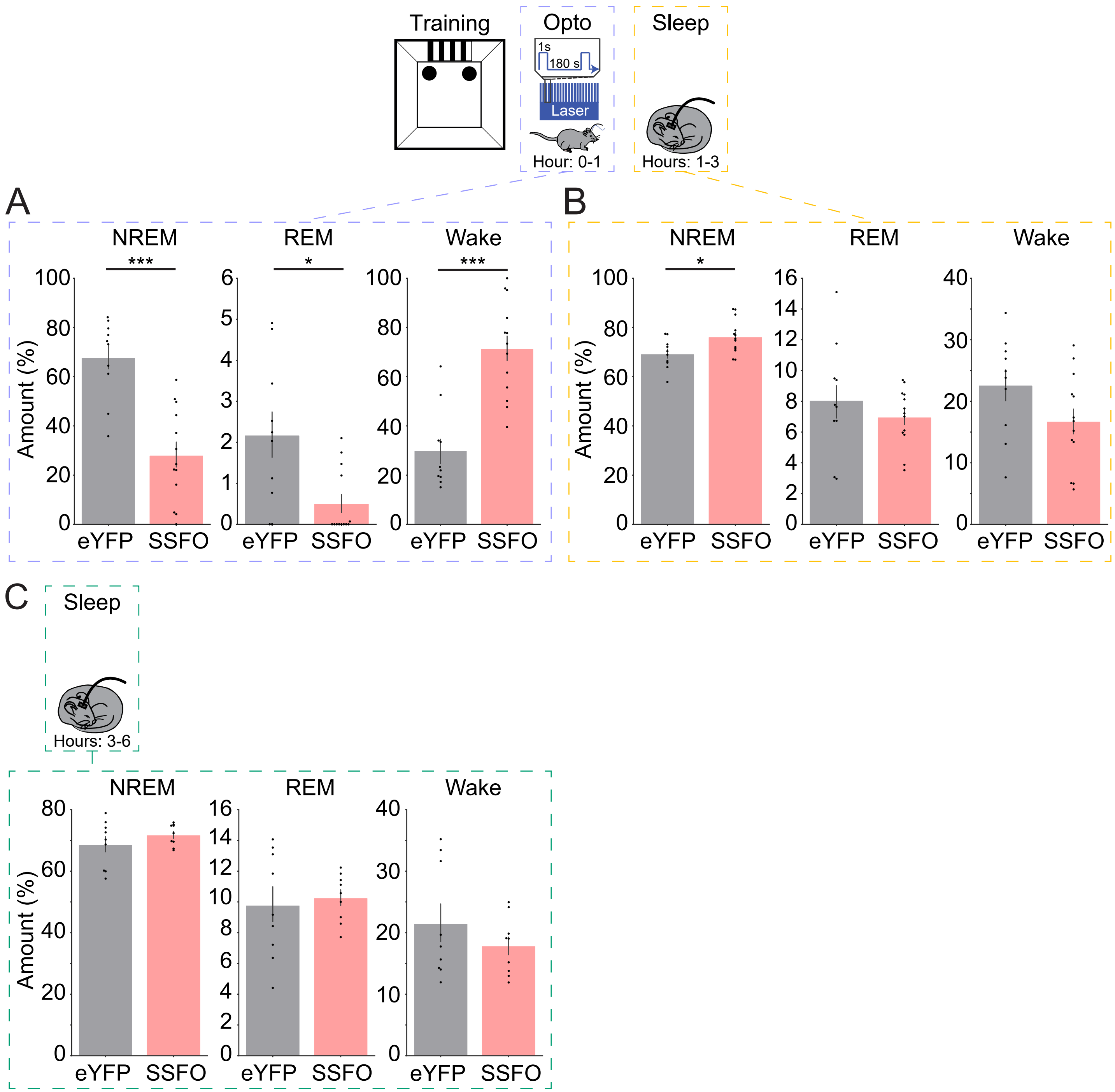

Training
Opto
1s
180 s
Laser
Hour: 0-1
Sleep
Hours: 1-3
A
NREM
REM
Wake
***
*
***
Amount (%)
eYFP
SSFO
B
NREM
REM
Wake
*
Amount (%)
eYFP
SSFO
C
Sleep
Hours: 3-6
NREM
REM
Wake
Amount (%)
eYFP
SSFO

Figure S2

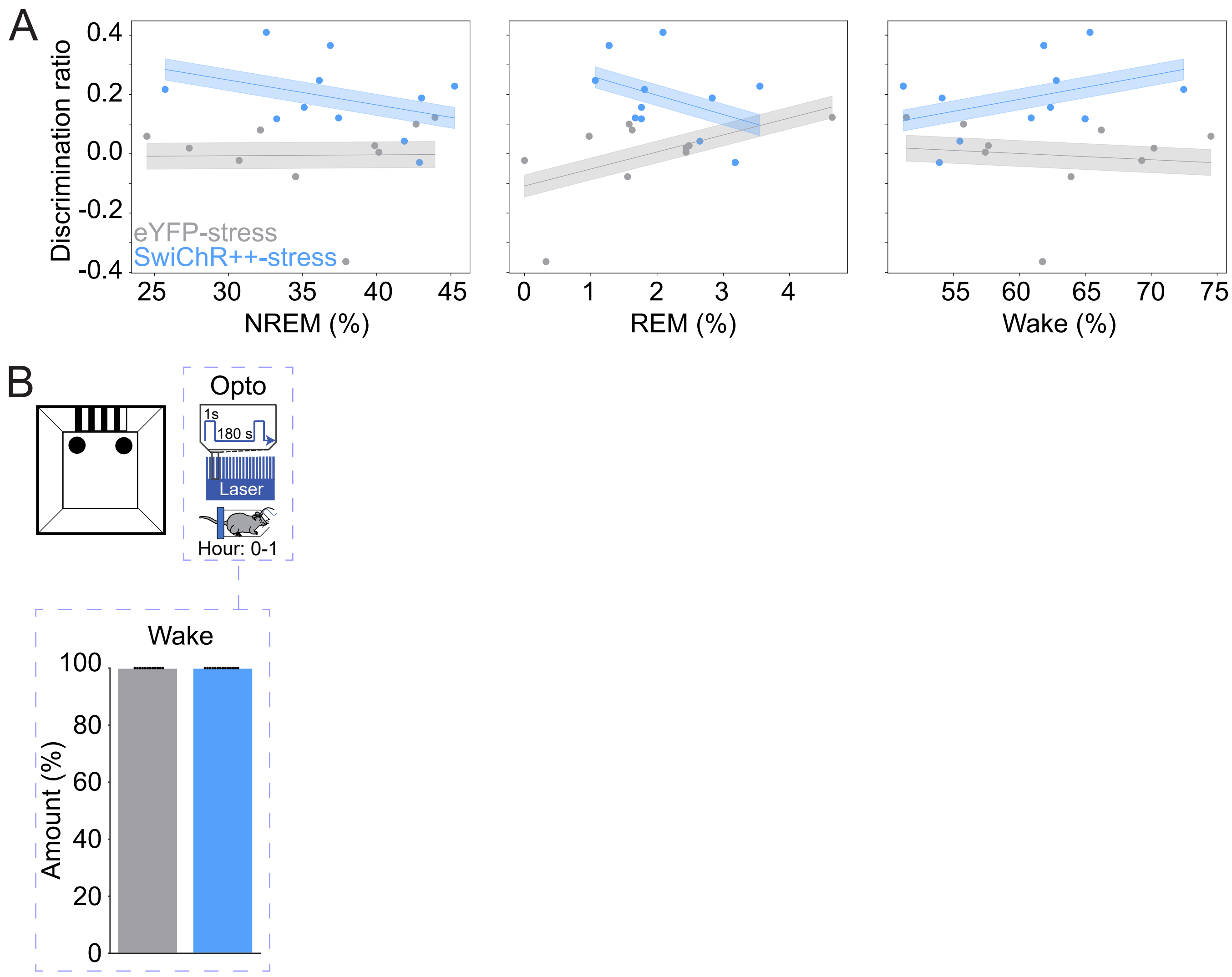
A
0.4
0.2
0.0
-0.2
-0.4
Discrimination ratio
eYFP-stress
SwiChR++-stress
25
30
35
40
45
NREM (%)
0
1
2
3
4
REM (%)
55
60
65
70
75
Wake (%)
B
Opto
1s
180 s
Laser
Hour: 0-1
Wake
100
80
60
40
20
0
Amount (%)

Figure S3

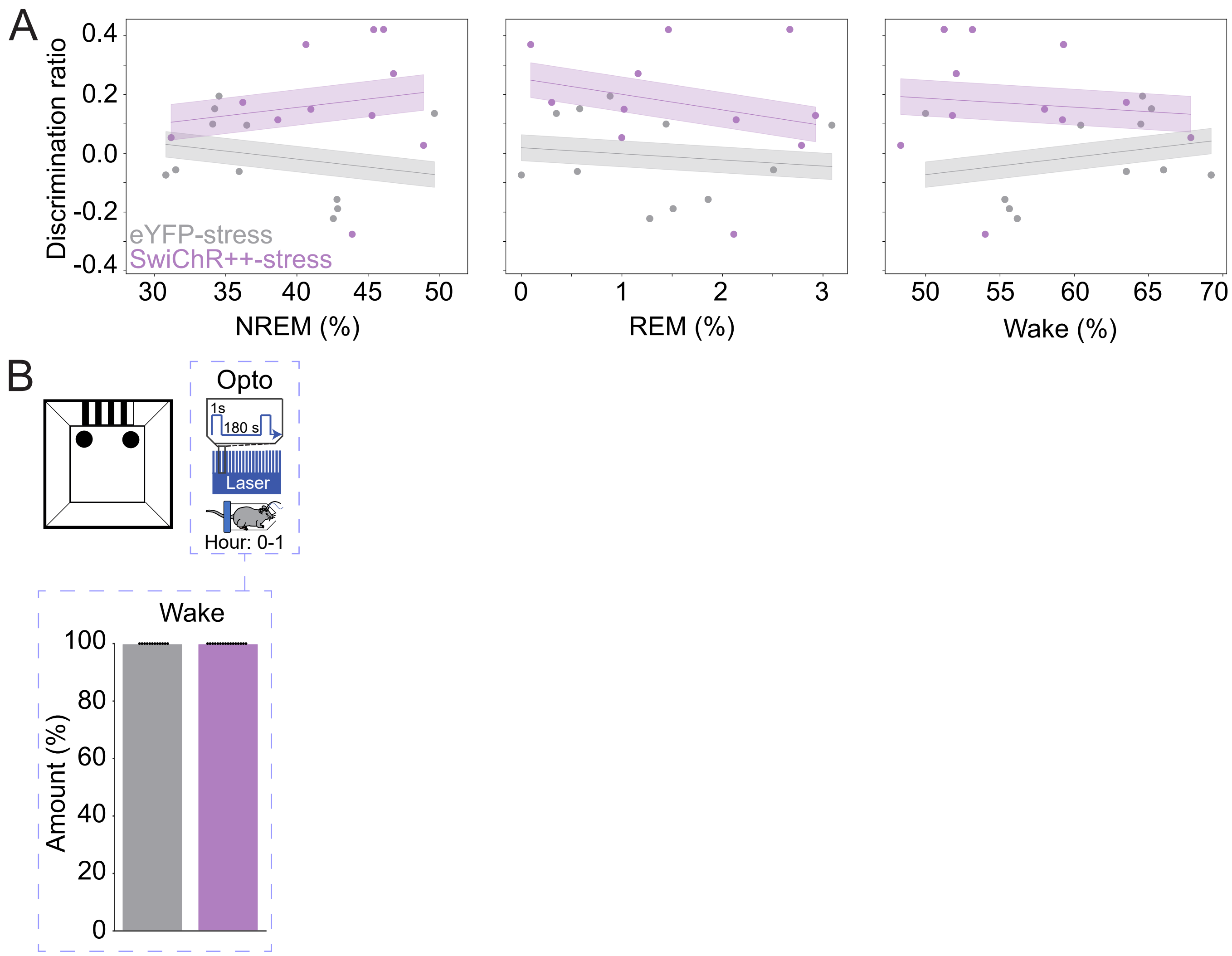
A
Discrimination ratio
0.4
0.2
0.0
-0.2
-0.4
eYFP-stress
SwiChR++-stress
30
35
40
45
50
NREM (%)
0
1
2
3
REM (%)
50
55
60
65
70
Wake (%)
B
Opto
1s
180 s
Laser
Hour: 0-1
Wake
100
80
60
40
20
0
Amount (%)

| Figure | Statistical test | Values | Sample size |
|---|---|---|---|
| 1D | Paired t-test | Test vs. Training<br>eYFP: T(8) = 3.032, P = 0.0163<br>SSFO: T(9) = 0.1356, P = 0.8951 | 9 eYFP mice<br>10 SSFO mice |
| 1E | T-test | SSFO vs. eYFP<br>T(17) = 3.087, P = 0.0067 | 9 eYFP mice<br>10 SSFO mice |
| 1G | T-test | eYFP vs. SSFO<br>NREM: T(14.594) = 2.894, P = 0.011<br>REM: T(13.012) = 1.445, P = 0.172<br>Wake: T(15.109) = -2.900, P = 0.011 | 10 eYFP mice<br>13 SSFO mice |
| 1H | Linear Mixed Model | NREM: Coef = 0.009, Std.Err = 0.006, z = 1.572, P = 0.116<br>Group: Coef = -0.152, Std. Err = 0.253, z = -0.603, P = 0.547<br><br>REM: Coef = 0.024, Std.Err = 0.020, z = 1.202, P = 0.230<br>Group: Coef = -0.203, Std. Err = 0.255, z = -0.794, P = 0.427<br><br>Wake: Coef = -0.009, Std.Err = 0.005, z = -1.716, P = 0.086<br>Group: Coef = -0.143, Std. Err = 0.250, z = -0.572, P = 0.567 | 9 eYFP mice<br>10 SSFO mice |
| 2C | Paired t-test | Test vs. Training<br>Control: T(11) =2.493, P = 0.0299<br>Stress: T(8) = 0.5953, P = 0.5681 | 12 control mice<br>9 stress mice |
| 2D | T-test | Stress vs. Control<br>T(19) = 2.133, P = 0.0462 | 12 control mice<br>9 stress mice |
| 2H | Paired t-test | Test vs. Training<br>eYFP-stress: T(10) = 0.2191, P = 0.8309<br>SwiChR++-stress: T(10) = 5.342, P = 0.0003 | 11 eYFP-stress mice<br>11 SwiChR++-stress mice |
| 2I | T-test | SwiChR++-stress vs. eYFP-stress<br>T(20) = 3.352, P = 0.0032 | 11 eYFP-stress mice<br>11 SwiChR++-stress mice |
| 2K | T-test | eYFP-stress vs. SwiChR++-stress<br>NREM: T(23.311) = -1.775, P = 0.089<br>REM: T(18.324) = -1.263, P = 0.222<br>Wake: T(23.108) = 1.829, P = 0.080 | 12 eYFP-stress mice<br>14 SwiChR++-stress mice |
| 3D | Paired t-test | Test vs. Training<br>eYFP: T(9) = 3.213, P = 0.0106<br>SwiChR++: T(9) = 2.291, P = 0.0477 | 10 eYFP mice<br>10 SwiChR++ mice |
| 3E | T-test | SwiChR++ vs. eYFP<br>T(18) = 0.9133, P = 0.3731 | 10 eYFP mice<br>10 SwiChR++ mice |
| 3G | T-test | eYFP vs. SwiChR++<br>NREM: T(24.868) = 0.628, P = 0.536<br>REM: T(18.307) = 0.565, P = 0.579<br>Wake: T(24.130) = -0.702, P = 0.490 | 13 eYFP mice<br>15 SwiChR++ mice |
| 4C | T-test | Stress vs. Control<br>T(4) = 9.725, P = 0.0006 | 3 control mice<br>3 stress mice<br>(averaged 3 sections/mouse, bilateral) |
| 4F | T-test | SSFO vs. eYFP<br>T(7) = 2.762, P = 0.0280 | 4 eYFP mice<br>5 SSFO mice<br>(averaged 2 sections/mouse, unilateral) |
| 4J | Paired t-test | Test vs. Training<br>eYFP-stress: T(10) = 0.2768, P = 0.7876<br>SwiChR++-stress: T(10) = 2.974, P = 0.0139 | 11 eYFP-stress mice<br>11 SwiChR++-stress mice |
| 4K | T-test | SwiChR++-stress vs. eYFP-stress<br>T(20) = 2.320, P = 0.0310 | 11 eYFP-stress mice<br>11 SwiChR++-stress mice |
| 4M | T-test | eYFP-stress vs. SwiChR++-stress<br>NREM: T(20.024) = -2.381, P = 0.027<br>REM: T(24.157) = -0.664, P = 0.513<br>Wake: T(21.458) = 2.405, P = 0.025 | 12 eYFP-stress mice<br>16 SwiChR++-stress mice |
| S1A | T-test | eYFP vs. SSFO<br>NREM: T(20.737) = 5.354, P = 0.000027<br>REM: T(11.830) = 2.766, P = 0.017<br>Wake: T(20.798) = -5.499, P = 0.000019 | 10 eYFP mice<br>13 SSFO mice |
| S1B | T-test | eYFP vs. SSFO<br>NREM: T(20.591) = -2.546, P = 0.019<br>REM: T(12.551) = 0.851, P = 0.411<br>Wake: T(18.834) = 1.761, P = 0.094 | 10 eYFP mice<br>13 SSFO mice |
| S1C | T-test | eYFP vs. SSFO<br>NREM: T(16) = -1.109, P = 0.284<br>REM: T(16) = -0.381, P = 0.708<br>Wake: T(16) = 1.046, P = 0.311 | 9 eYFP mice<br>9 SSFO mice |
| S2A | Linear Mixed Model | NREM: Coef = -0.004, Std.Err = 0.005, z = -0.724, P = 0.469<br>Group: Coef = 0.200, Std. Err = 0.201, z = 0.994, P = 0.320<br><br>REM: Coef = 0.022, Std.Err = 0.029, z = 0.765, P = 0.444<br>Group: Coef = 0.185, Std. Err = 0.200, z = 0.923, P = 0.356<br><br>Wake: Coef = 0.002, Std.Err = 0.005, z = 0.538, P = 0.591<br>Group: Coef = 0.198, Std. Err = 0.202, z = 0.981, P = 0.326 | 10 eYFP-stress mice<br>11 SwiChR++-stress mice |
| S2B | T-test | Wake: All mice recorded had 100% wake; variance = 0,<br>preventing standard t-test calculation. | 12 eYFP-stress mice<br>14 SwiChR++-stress mice |
| S3A | Linear Mixed Model | NREM: Coef = -0.000, Std.Err = 0.007, z = -0.065, P = 0.948<br>Group: Coef = 0.178, Std. Err = 0.272, z = 0.656, P = 0.512<br><br>REM: Coef = -0.038, Std.Err = 0.041, z = -0.909, P = 0.363<br>Group: Coef = 0.189, Std. Err = 0.265, z = 0.713, P = 0.476<br><br>Wake: Coef = 0.001, Std.Err = 0.007, z = 0.207, P = 0.836<br>Group: Coef = 0.183, Std. Err = 0.135, z = 1.355, P = 0.175 | 11 eYFP-stress mice<br>11 SwiChR++-stress mice |
| S3B | T-test | Wake: All mice recorded had 100% wake; variance = 0,<br>preventing standard t-test calculation. | 12 eYFP-stress mice<br>16 SwiChR++-stress mice |

**Table S1**
